# Robustness encoded across essential and accessory replicons in an ecologically versatile bacterium

**DOI:** 10.1101/209916

**Authors:** George C diCenzo, Alex B Benedict, Marco Fondi, Graham C Walker, Turlough M Finan, Alessio Mengoni, Joel S Griffitts

**Affiliations:** Department of Biology, University of Florence, Sesto Fiorentino, FI, 50019, Italy.; Department of Microbiology and Molecular Biology, Brigham Young University, Provo, UT, 84602, USA.; Department of Biology, Massachusetts Institute of Technology, Cambridge, MA, 02139, USA.; Department of Biology, McMaster University, Hamilton, ON, L8S 4K1, Canada.

**Keywords:** genetic interactions, synthetic lethality, systems biology, Tn-seq, constraint based modelling

## Abstract

Bacterial genome evolution is characterized by gains, losses, and rearrangements of functional genetic segments. The extent to which genotype-phenotype relationships are influenced by large-scale genomic alterations has not been investigated in a high-throughput manner. In the symbiotic soil bacterium *Sinorhizobium meliloti,* the genome is composed of a chromosome and two large extrachromosomal replicons (pSymA and pSymB, which together constitute 45% of the genome). Massively parallel transposon insertion sequencing (Tn-seq) was employed to evaluate contributions of chromosomal genes to fitness in both the presence and absence of these extrachromosomal replicons. Ten percent of chromosomal genes from diverse functional categories are shown to genetically interact with pSymA and pSymB. These results demonstrate the pervasive robustness provided by the extrachromosomal replicons, which is further supported by constraint-based metabolic modelling. A comprehensive picture of core *S. meliloti* metabolism was generated through a Tn-seq-guided *in silico* metabolic network reconstruction, producing a core network encompassing 726 genes. This integrated approach facilitated functional assignments for previously uncharacterized genes, while also revealing that Tn-seq alone misses over a quarter of wild type metabolism. This work highlights the strong functional dependencies and epistatic relationships that may arise between bacterial replicons and across a genome, while also demonstrating how Tn-seq and metabolic modelling can be used together to yield insights not obtainable by either method alone.

## Introduction

The prediction of genotype-phenotype relationships is a fundamental goal of genetic, biomedical, and eco-evolutionary research, and this problem underpins the design of synthetic microbial systems for biotechnological applications [1]. The last decades have witnessed a shift away from the functional characterization of single genes towards whole-genome, systems-level analyses [for recent reviews, see [2,3]]. Such studies have been facilitated by the development of methods that allow for the direct interrogation of a genome to determine all genetic elements required for adaptation to a specified environment. Two primary methods are *in silico* metabolic modelling [4,5], and massively parallel sequencing of transposon insertions in bacterial mutant libraries (Tn-seq) [6,7].

The process of *in silico* genome-scale metabolic modelling consists of two stages. First, a reconstruction of all cellular metabolism is built that contains all reactions expected to be present, as well as which genes encode the enzymes performing each reaction, thereby linking genetics to metabolism [8]. Next, mathematical models such as flux balance analysis (FBA) are used to simulate the flux distribution through the reconstructed metabolic network [9], which can be used to predict how environmental perturbations or gene disruptions influence growth phenotypes. This approach allows for phenotypic predictions of all possible single, double, or higher-order gene deletion mutations within a matter of days [10,11], something that is infeasible using a direct experimental approach. However, the quality of the predictions is highly dependent on the accuracy of the metabolic reconstruction. Outside of a few model species like *Escherichia coli,* experimental genetic and biochemical data are not available at the resolution necessary to provide accurate assignment of all metabolic gene functions.

The Tn-seq approach involves the generation of a library of hundreds of thousands of mutant clones, each containing a single transposon insertion at a random genomic location [12]. The library of pooled clones is then cultured in the presence of a defined environmental challenge. Insertions resulting in altered fitness in the environment under investigation become under‐ or over-represented in the population, and this is monitored by deep sequencing to identify the genomic location and frequency of all transposon insertions. This approach is imperfect, as important biochemical functions may be encoded redundantly in the genome [13–15], and the loss of some essential genes can be compensated for by evolution of alternative cellular processes [16]. Moreover, fitness changes brought about by mutation in one gene may be dependent on mutation of a second gene bearing no resemblance to the first—a phenomenon known as a genetic interaction [17,18]. Such genetic interactions may cause the apparent functions of some genes to be strictly dependent on their genomic environment [19]. In other words, a gene may be essential for growth in one organism, but its orthologous counterpart in another organism may be non-essential. This significantly complicates efforts to generalize genotype-phenotype relationships [20].

Resolving the problem of genome-conditioned gene function is of broad significance in the areas of functional genomics, population genetics, and synthetic biology. For example, the ability to design and build optimized minimal cell factories on the basis of single-mutant fitness data is expected to present numerous complications [21], as evidenced by the recent effort to rationally build a functional minimal genome [22]. Tn-seq studies have suggested there is as little as 50% to 25% overlap in the essential genome of any two species [23–25]. As a striking example, 210 of the Tn-seq determined essential genes of *Pseudomonas aeruginosa* PA14 are not even present in the genome of *P. aeruginosa* PAO1 [26]. Comparison of Tn-seq data for *Shigella flexneri* with the deletion analysis data for closely related *E. coli* suggested only a small number of genes were specifically essential in one species. Mutation of about 100 genes, however, appeared to result in a growth rate decrease specifically in *E. coli* [27]. Similarly, comparison of Tn-seq datasets from two *Salmonella* species revealed that mutation of nearly 40 genes had a stronger growth phenotype in one of the two species [28]. Overall, these studies suggest that the genomic environment (here defined as the genomic components that may vary from organism to organism) influences the fitness contributions of a significant proportion of an organism’s genes. However, no large-scale analysis has been performed that directly illustrates how the phenotypes of individual genes are impacted when a small or large part of the genome is modified.

Here, we provide a quantitative, genome-scale evaluation of how large-scale genomic variance influences genotype-phenotype relationships. We have accomplished this in a way that minimizes the effects of laboratory-to-laboratory variation, and removes the effects of complex genome evolution. The model system used is *Sinorhizobium meliloti,* an α-proteobacterium whose 6.7-Mb genome consists of a chromosome and two additional replicons, the pSymA megaplasmid and the pSymB chromid. The pSymA and pSymB replicons constitute 45% of the *S. meliloti* genome (∼2,900 genes); yet, by simply transferring only two essential genes from pSymB to the chromosome, both pSymA and pSymB can be completely removed from the genome, yielding a viable single-replicon organism [29]. We report a comparison of gene essentiality (via Tn-seq) for wild-type *S. meliloti* and the single-replicon derivative. This analysis was supplemented by an *in silico* double gene deletion analysis of a *S. meliloti* genome-scale metabolic network reconstruction. We further examine how integration of Tn-seq data with *in silico* metabolic modelling, through a Tn-seq-guided reconstruction process, overcomes the limitations of using either of these approaches in isolation to develop a consolidated view of the core metabolism of the organism. This process produced a fully referenced *core S. meliloti* metabolic reconstruction.

## RESULTS

### Development and validation of the Tn5-based transposon Tn5-714

In order to interrogate the *S. meliloti* genome using a Tn-seq based approach, we first developed a new construct based on the Tn5 transposon as described in the Materials and Methods. The resulting transposon (Figure S1) contains constitutive promoters reading out from both ends of the transposon to ensure the production of non-polar mutations. Analysis of the insertion site locations validated that the transposon performed largely as expected. Gene disruptions caused by transposon insertions were confirmed to be non-polar as illustrated by the case reported in Figure 1, and there was no strong bias in the distribution of insertions around the chromosome (Figures 2A, S2). However, there did appear to be somewhat of a bias for integration of the transposon in GC rich regions (Figure S3). Given the high GC content (62.7%) of the *S. meliloti* chromosome, it is unlikely that this moderate bias had a discernable influence on the results of this study.

**Figure 1.**
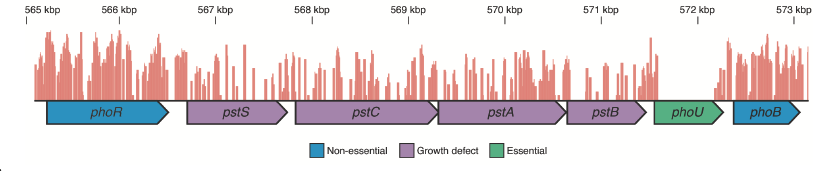
Visualization of the location of transposon insertion sites. An image of the *pst* locus of *S. meliloti* generated using the Integrative Genomics Viewer [85]. Chromosomal nucleotide positions are indicated along the top of the image, and the location of transposon insertions are indicated by the red bars. Non-essential genes contain a high density of transposon insertions, whereas essential genes have few to no transposon insertions. Genes are color coded based on their fitness classification, and transcripts are indicated by the arrows below the genes. The *pstS, pstC, pstA, pstB, phoU,* and *phoB* genes are co-transcribed as a single operon [86], and previous work demonstrated that polar *phoU* mutations are lethal in *S. meliloti,* whereas non-polar mutations are not lethal [87]. The lack of insertions within the *phoU* coding region is therefore consistent with the non-polar nature of the transposon.

**Figure 2.**
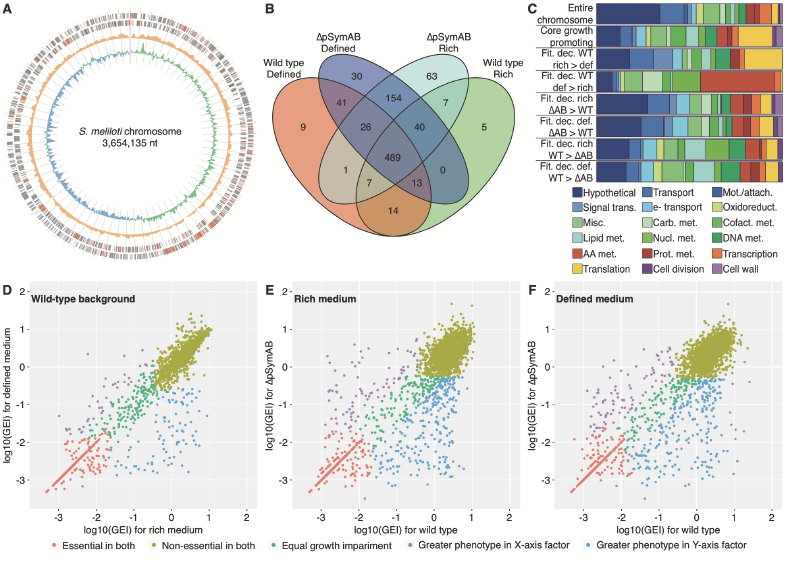
Characteristics of the core genetic components of *S. meliloti.* (A) A plot of the *S. meliloti* chromosome is shown. From the outside to inside: positive strand coding regions, negative strand coding regions, total insertion density, and GC skew. For the positive and negative strands, red lines indicate the core 489 growth promoting genes. The insertion density displays the total transposon insertions across all experiments over a 10,000-bp window. The GC skew was calculated over a 10,000-bp window, with green showing a positive skew and blue showing a negative skew. Tick marks are every 50,000 bp. (B) A comparison of the overlap between the growth promoting genome (Group I and II genes) of each Tn-seq data set. Each data set is labelled with the strain (wild type or ΔpSymAB) and the growth medium (defined medium or rich medium). (C) Functional enrichment plots for the indicated gene sets. Name abbreviations: Fit - fitness; Dec - decrease; WT - wild type; AAB - ΔpSymAB; Def - defined medium; Rich - rich medium. For example, ‘Fit. dec. WT def > rich’ means the genes with a greater fitness decrease in wild type grown in defined medium compared to rich medium. Legend abbreviations: AA - amino acid; Attach - attachment; Carb - carbohydrate; Cofact - cofactor; e— electron; Met - metabolism; Misc - miscellaneous; Mot - motility; Nucl - nucleotide; Oxidoreduct - oxidoreductase activity; Prot - protein; Trans - transduction. (D-F) Scatter plots comparing the fitness phenotypes, shown as the logj_0_ of the GEI scores (Gene Essentiality Index scores; i.e., number of insertions within the gene divided by gene length in nucleotides) of (D) wild type grown in rich medium versus wild type grown in defined medium, (E) wild type grown in rich medium versus ΔpSymAB grown in rich medium, and (F) wild type grown in defined medium versus ΔpSymAB grown in defined medium.

### Overview of the Tn-seq output

The Tn-seq experiments reported here were undertaken with two primary aims: i) to identify the core set of genes contributing to *S. meliloti* growth in laboratory conditions, and ii) to determine the extent to which the phenotypic consequence of a gene deletion is influenced by the genomic environment (i.e. presence/absence of the secondary replicons). To accomplish this, Tn-seq libraries of two *S. meliloti* strains were prepared: a wild type strain (designated RmP3499) containing the entire genome, and a strain with both the pSymA and pSymB replicons removed (designated RmP3496 or ΔpSymAB; strains described previously in [30]). Transposon library sizes were skewed to compensate for the difference in genome sizes, resulting in nearly identical insertion site density for each library (Table S1). Both libraries were passed through selective growth regimens in either complex BRM broth (rich medium) or minimal VMM broth (defined medium) in duplicates. Following approximately nine generations of growth, the location of the transposon insertions in the population was determined, a gene essentiality index (GEI) was calculated for all chromosomal genes, and each gene was classified into one of five fitness categories (Table 1) using the procedure described in the Materials and Methods. Four genes *(pdxJ, fumC, smc01011, smc03995),* including two of unknown function, were independently mutated in the wild-type background, and in all cases, the mutations yielded the expected no-growth phenotype (Figure S4), supporting the accuracy of the Tn-seq output. All Tn-seq data is available as Data Set S1.

**Table 1.** Fitness classification of chromosomal genes. Genes were ranked from lowest to highest GEI, with the lowest GEI being at the 0 percentile and the highest GEI being at the 100^th^ percentile. The approximate break points for the groupings, determined as described in the Materials and Methods, are shown for each condition.

| Group | Description | GEI percentile range |  |  |  |
| --- | --- | --- | --- | --- | --- |
| | | Wild type, rich medium | ΔpSymAB, rich medium | Wild type, defined medium | ΔpSymAB, defined medium |
| I | Essential | 0-12 | 0-14 | 0-12 | 0-14 |
| II | Strong growth defect | 12-17 | 14-23 | 12-18 | 14-24 |
| III | Moderate growth defect | 17-36 | 23-49 | 18-28 | 24-47 |
| IV | Little to no growth impact | 36-100 | 49-96 | 28-99 | 47-99 |
| V | Growth improvement | N/A | 96-100 | 99-100 | 99-100 |

A strong correlation was observed between the number of insertions per gene in each set of duplicates (Figure S5), indicating that there was high reproducibility of the results and that differences between conditions were unlikely to reflect random fluctuations in the output. On average, insertions were found in 190,000 unique chromosomal positions with a median of 39 unique insertion positions per gene (Table S1). The similarity in the number of unique insertion positions between samples suggested that differences in the Tn-seq outputs were also unlikely to be an artefact of the quality of the libraries.

### Elucidation of the core genetic components of *S. meliloti.*

There were 307 genes classified as essential independently of growth medium or strain (Figure S6). This set of 307 genes includes those encoding functions commonly understood to be essential: the DNA replication apparatus, the four RNA polymerase subunits, the housekeeping sigma factor, the general transcriptional termination factor Rho, 40 out of 55 of the annotated ribosomal protein subunits, 18 out of 20 of the annotated aminoacyl-tRNA synthetases, and 6 out of 10 of the annotated ATP synthase subunits. Considering genes classified as essential plus those genes whose mutation resulted in a large growth defect (Groups I and II in Table 1), a core growth promoting genome of 489 genes, representing ∼ 15% of the chromosome, was identified (Figure 2B). This expanded list includes 51 out of 55 of the annotated ribosomal protein subunits, 19 out of 20 of the annotated aminoacyl-tRNA synthetases, and 9 out of 10 of the annotated ATP synthase subunits These 489 genes appeared to be mostly dispersed around the chromosome, although there was a bias for these genes to be found in the leading strand (Figure 2A). Based on published RNA-seq data for *S. meliloti* grown in a glucose minimal medium, these 489 genes tend to be highly expressed, with a median expression level above the 90% percentile (Figure S7). Compared to the entire chromosome (Fisher exact test, p-value < 0.05 following a Bonferroni correction for 18 tests), this set of 489 genes was enriched for genes involved in translation (5.2-fold), lipid metabolism (2.7-fold), cofactor metabolism (3.3-fold), and electron transport (2.1-fold), whereas genes involved in transport (2.1-fold), motility/attachment (9.4-fold), and hypothetical genes (2.7-fold) were under-represented (Figure 2C). Additionally, cell wall (2.2-fold) and cell division (2.3-fold) were over-represented while transcription (1.9-fold) was under-represented (Figure 2C), although these differences where not considered statistically significant.

A clear influence of the growth medium on the fitness phenotypes of gene mutations was observed. The degree to which mutant phenotypes were impacted by growth medium type is reflected in the synthetic medium index (SMI) calculated as described in the Materials and Methods. Focusing on the wild-type strain, a core of 519 genes were identified as contributing equally to growth in both media (Figure 2D). Forty genes were identified as more important during growth in rich medium than in defined medium, and these genes had a median SMI score of 7 (values of 1 and −1 are neutral). Only translation functions (5.8-fold) displayed a statistically significant enrichment in these genes, which may reflect the faster growth rate in the rich medium (Figure S8), while there was also a non-statistically significant enrichment in signal transduction (5.1-fold) (Figure 2C). The extent of specialization for growth in the defined medium was more pronounced; 93 genes were more important during growth in the defined medium with a median SMI score of −20. These genes were enriched (statistically significant) in amino acid (9.0-fold) and nucleotide (6.7-fold) metabolism presumably due to the requirement of their biosynthesis, and carbohydrate metabolism (3.6-fold) likely as the sole carbon source was a carbohydrate (Figure 2C). The same overall pattern was observed between media for the ΔpSymAB strain (Figure S9).

### Mutant fitness phenotypes are strongly influenced by their genomic environment

The Tn-seq data sets for the wild-type and the ΔpSymAB strains were compared to evaluate the robustness of the observed fitness phenotypes in response to changes in the gene’s genomic environment. Similar results were observed for both growth media, suggesting that the results were generalizable and not medium specific. Depending on the medium, either 484 or 488 genes had an equal contribution to growth in both strains, 81 or 89 genes led to stronger growth impairment when mutated in wild-type cells, and either 250 or 251 genes led to stronger growth impairment when mutated in ΔpSymAB cells (Figures 2E, 2F, and Table 2). Only minor functional bias was observed in the genes that displayed larger fitness defects in the ΔpSymAB background (Figure 2C); in both media, only electron transport (3-fold) and oxidoreductases (9.5-fold) were over-and under-represented, respectively. Similarly, little functional bias was detected in genes with larger fitness defects in the wild-type background (Figure 2C); in both media, lipid metabolism (4.5-fold) and hypothetical genes (2-fold) were over-and under-represented, respectively, while nucleotide metabolism (5.5-fold) was also enriched in the rich medium. Overall, these results were consistent with pervasive effects of the genomic environment on the genotype-phenotype relationship that was largely independent of the biological role of the gene products.

**Table 2.**
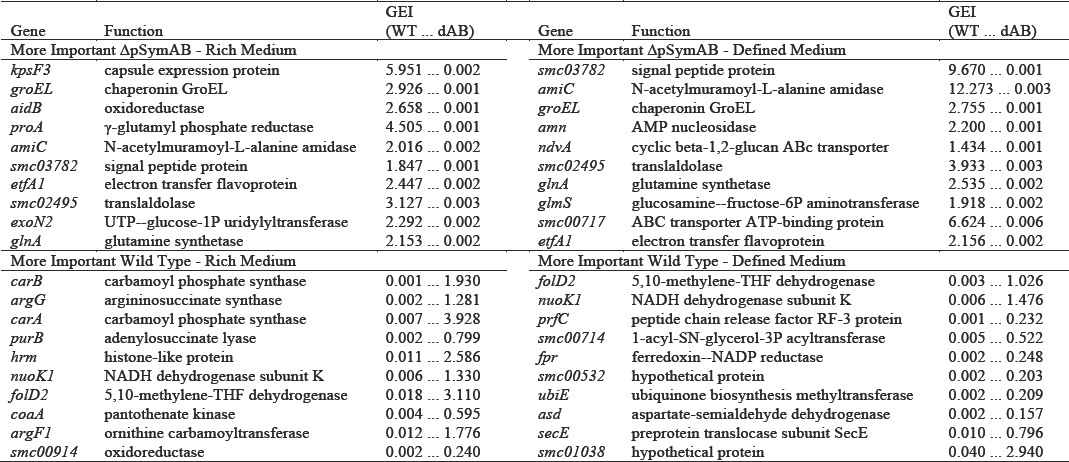
Sample genes showing strain specific phenotypes. The top ten genes from each of the indicated groupings, as determined based on the ratio of GEI scores of the two strains, are shown. GEI (Gene Essentiality Index) scores are shown first for the wild type (WT) followed by the scores for the ΔpSymAB (dAB) strain.

Approximately half (9 of 16) of the genes that were independently mutated in both strains yielded the expected phenotypes on rich agar plates (Figures S10). Of the other seven genes, which were expected to be essential specifically in the ΔpSymAB strain, at least three were non-lethal but displayed obvious growth rate defects or extended lag phases during liquid culture experiments (Table S2 and Figure S11). The remaining three genes may represent false positives from the Tn-seq screen, or may reflect differences in the growth conditions, namely, competitive growth versus isogenic growth. Nevertheless, the observation that at least 75% of the selected genes were confirmed to have a genome content-dependent fitness phenotype validates that the large majority of the strain specific phenotypes observed in the Tn-seq screen represent true differences.

### Level of genetic and phenotypic conservation of the essential *S. meliloti* genes

Several recent studies have used Tn-seq to study the essential genome of *Rhizobium leguminosarum* [31–33]. We compared our Tn-seq datasets with those reported in by Perry *et al* [32] to examine the conservation of the essential genome of these two closely related N_2_-fixing species. Putative orthologs for ∼ 75% of all *S. meliloti* chromosomal genes were identified in *R. leguminosarum* via a Blast Bidirectional Best Hit (Blast-BBH) approach (Data Set S2). Much higher conservation of the growth promoting genome was observed; 97% of the 489 core growth promoting genes and 99% of the 307 core essential genes had a putative ortholog in *R. leguminosarum.* However, conservation of the gene did not necessarily correspond to conservation of the phenotype. Considering only the 303 conserved core essential *S. meliloti* genes (as these were the least likely to have been falsely identified as essential), 8% (25 of 303) of their orthologous genes were classified as having little contribution to growth on defined medium in *R. leguminosarum* (Figure 3A). An additional 34 genes were considered to be non-essential but growth defective when mutated (Figure 3A). Independent mutation of two genes (*fumC*, *pdxJ)* identified as specifically essential in *S. meliloti* confirmed their essentiality (Figure S4), supporting the Tn-seq data. A similar pattern is observed starting with the *R. leguminosarum* genes classified as essential in both minimal and complex medium by Perry *et al.* [32]. Of the 241 core essential *R. leguminosarum* genes with an ortholog on the *S. meliloti* chromosome, 21 (9%) of the orthologs were classified as non-essential in *S. meliloti* for growth in defined medium, while an additional 8 were considered to have a moderate growth defect (Figure 3B).

**Figure 3.**
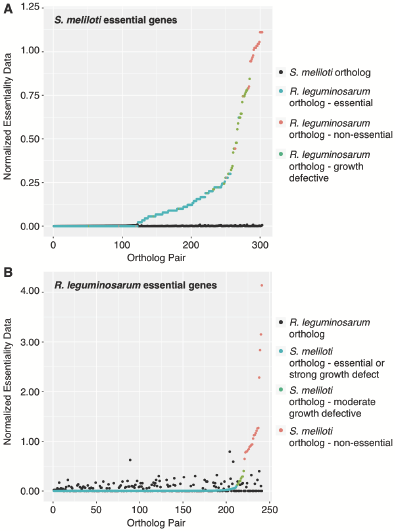
Comparison of *S. meliloti* and *R. leguminosarum* Tn-seq data. (A) The fitness phenotypes of essential *S. meliloti* genes, as determined in this study, is compared to the fitness phenotypes of the orthologous *R. leguminosarum* genes, as determined by Perry *et al.* [32]. *S. meliloti* orthologs are shown in black, while the *R. leguminosarum* orthologs are colored according to their classification by Perry *et al.* [32]. (B) The fitness phenotypes of essential *R. leguminosarum* genes is compared to the fitness phenotypes of the orthologous *S. meliloti* genes. *R. leguminosarum* orthologs are shown in black, while the *S. meliloti* orthologs are colored according to their classification in this study. (A,B) Normalized fitness values are used to facilitate direct comparison between the studies as different output statistics were calculated. For *S. meliloti*, the GEI score of each gene for wild type grown in minimal medium broth was divided by the median GEI for all genes under the same conditions. For *R. leguminosarum,* the insertion density of each gene during growth on minimal medium plates was divided by the median insertion density of all strains.

To further test the species specificity of the above-mentioned genes, the experiment was replicated *in silico.* Fifteen of the 25 orthologs specifically essential in *S. meliloti* were present both in our existing *S. meliloti* genome-scale metabolic model [34] as well as in a draft *R. leguminosarum* metabolic model (see Materials and Methods). Flux balance analysis was used to examine the *in silico* effect of deleting these 15 pairs of orthologs on growth. Three pairs of orthologs were classified as essential in both models, five were classified as non-essential in both models, and seven were classified as essential specifically in the *S. meliloti* model. Thus, at least half of the gene essentiality differences observed in the Tn-seq data are corroborated by the *in silico* metabolic simulation, despite the preliminary nature of the draft *R. leguminosarum* model. An *in silico* analysis of the genes identified as specifically essential in *R. leguminosarum* on the basis of the Tn-seq data was not performed as only two of these genes were present in the *R. leguminosarum* model.

### *In silico* analyses support a high potential for genetic redundancy in the *S. meliloti* genome

The results of the previous two sections are consistent with a strong genomic environment effect on the phenotypic consequences of gene mutations. One possible explanation is the presence of widespread genetic redundancy, at the gene and/or pathway level. In support of this, ∼ 14% of chromosomal genes had a Blast-BBH hit when the chromosomal proteome was compared against the combined pSymA/pSymB proteome (Data Set S3). Therefore, this phenomenon was further explored using a constraint-based metabolic modelling approach.

We first tested the *in silico* effect of chromosomal single gene deletions on growth rate in the presence and absence of pSymA/pSymB (Figure 4A). This analysis identified 67 genes (∼ 7% of all chromosomal model genes) as having a more severely impaired growth phenotype when deleted in the absence of pSymA/pSymB genes, 38 of which were lethal. This appeared to be due to a combination of direct functional redundancy of the gene products as well as through metabolic bypasses, as deletion of 50 reactions dependent on chromosomal genes had a more severe phenotype in the absence of pSymA/pSymB, 42 of which were lethal (Figure S12).

**Figure 4.**
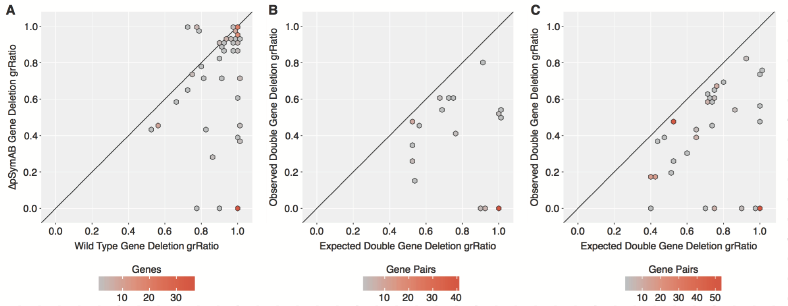
*In silico* analysis of genetic redundancy in *S. meliloti.* The effects of single or double gene deletion mutants were predicted *in silico* with the genome-scale *S. meliloti* metabolic model. (A) A scatter plot comparing the grRatio (growth rate of mutant / growth rate of non-mutant) for gene deletion mutations in the presence (wild type) versus absence (ΔpSymAB) of the pSymA/pSymB model genes. Genes whose deletion had either no effect or were lethal in both cases are not shown. (B) A scatter plot comparing the grRatio for each double gene deletion pair (where one gene was on the chromosome and the other on pSymA or pSymB) observed *in silico* versus the predicted grRatio based on the grRatio of the single deletions (grRatio1 * grRatio2). Only gene pairs with an observed grRatio at least 10% less than the expected are shown. (C) A scatter plot comparing the grRatio for each double gene deletion pair (both genes on the chromosome) observed *in silico* versus the predicted grRatio. Only gene pairs with an observed grRatio at least 10% less than the expected are shown. (A-C) The color of each hexagon is representative of the number of reactions plotted at that location, as illustrated by the density bar below each panel. The diagonal line serves as a reference line.

Next, a double gene deletion analysis was performed to examine the effect on growth rate of deleting every possible pair of model genes. This analysis suggested that 49 chromosomal genes had a more significant impact on growth than expected when simultaneously deleted with a single pSymA or pSymB gene (Figure 4B). Additionally, synthetic negative phenotypes were observed for 97 chromosomal genes when simultaneously deleted with another chromosomal gene (Figure 4C). Overall, 14% of chromosomal genes were predicted to have a synthetic negative phenotype when co-deleted with a second gene, consistent with a high potential for metabolic robustness being encoded by the *S. meliloti* genome, and with a significant influence of the genomic environment on the fitness phenotype of gene mutations.

### A consolidated view of core *S. meliloti* metabolism through Tn-seq-guided *in silico* metabolic reconstruction

The results described in the previous sections made it evident that a Tn-seq approach alone is insufficient to elucidate all processes contributing to growth in a particular environment. This is especially true if also considering non-essential metabolism that is nevertheless actively present in wild type cells, such as exopolysaccharide production. Moreover, it is difficult to fully comprehend the core functions of a cell by simply examining a list of essential genes and their predicted functions. We therefore attempted to overcome these limitations by using the Tn-seq data to guide a manual *in silico* reconstruction of the core metabolic processes of *S. meliloti.* A detailed description of this process is provided in the Materials and Methods. In brief, the existing metabolic model iGD1575 was treated as a database of reactions and gene-reaction associations. Each pathway involved in central carbon metabolism or the production of essential or non-essential biomass components (Table S3) were then rebuilt in a new (initially empty) reconstruction drawing from the reactions present in iGD1575. At the same time, the genes associated with each reaction were compared to the Tn-seq data and published literature to confirm the linkage of the correct gene(s) to each reaction.

The resulting model, termed iGD726 and included as in SBML format in File S2, is summarized in Figure 5 and Table 3, and the entire model including genes, reaction formulas, and references is provided as an easy to read Excel table in Data Set S4. The process of integrating the Tn-seq data with *in silico* metabolic reconstruction resulted in a major refinement of the core metabolism compared to the existing genome-scale model: 228 new reactions were added, 115 new genes were added, and the genes associated with 135 of the 432 reactions common to both reconstructions were updated. In addition to improving the metabolic reconstruction, this process significantly expanded the view of core *S. meliloti* metabolism compared to that gained solely through the application of Tn-seq. The genes associated with approximately one third of the iGD726 reactions were not detected as growth promoting in the Tn-seq datasets (Figure 5, Table 3). While many of the additional reactions present in iGD726 are due to the inclusion of non-essential biomass components, which are part of the wild type cell but are nonetheless dispensable for growth, others are from essential metabolic pathways (Figures 5, S13). Overall, the combined approach of integrating Tn-seq data and *in silico* metabolic modelling allowed for the development of a high-quality representation of core *S. meliloti* metabolism in a way that neither approach alone was capable of accomplishing.

**Figure 5.**
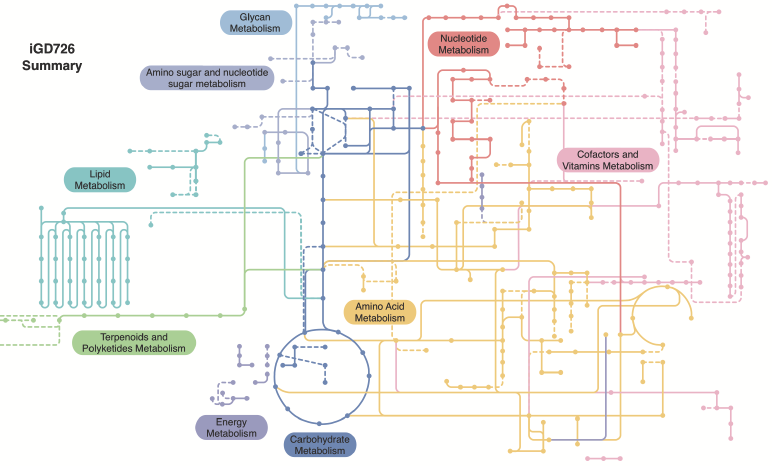
Summary schematic of core *S. meliloti* metabolism. The iGD726 core metabolic model was visualized using the iPath v2.0 webserver [74], which maps the reactions of the metabolic model to KEGG metabolic pathways; it therefore does not capture metabolism not present in the KEGG pathways included in iPath. Reactions and metabolites are colour coded according to their biological role, as indicated. Reactions whose associated genes were not identified as growth promoting in this study are in dashed lines.

**Table 3.**
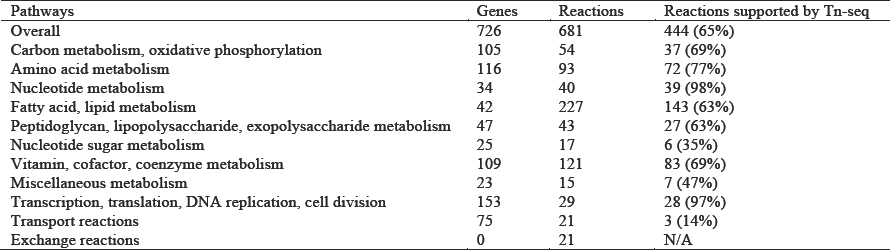
Summary of iGD726. The last column indicates reactions whose genes associations are supported by the Tn-seq data of this study. Percentage of all reactions in that category are indicated in brackets.

| Pathways | Genes | Reactions | Reactions supported by Tn-seq |
| --- | --- | --- | --- |
| Overall | 726 | 681 | 444 (65%) |
| Carbon metabolism, oxidative phosphorylation | 105 | 54 | 37 (69%) |
| Amino acid metabolism | 116 | 93 | 72 (77%) |
| Nucleotide metabolism | 34 | 40 | 39 (98%) |
| Fatty acid, lipid metabolism | 42 | 227 | 143 (63%) |
| Peptidoglycan, lipopolysaccharide, exopolysaccharide metabolism | 47 | 43 | 27 (63%) |
| Nucleotide sugar metabolism | 25 | 17 | 6 (35%) |
| Vitamin, cofactor, coenzyme metabolism | 109 | 121 | 83 (69%) |
| Miscellaneous metabolism | 23 | 15 | 7 (47%) |
| Transcription, translation, DNA replication, cell division | 153 | 29 | 28 (97%) |
| Transport reactions | 75 | 21 | 3 (14%) |
| Exchange reactions | 0 | 21 | N/A |

### Tn-seq-guided *in silico* metabolic reconstruction facilitates novel gene annotation

Over 20 of the reactions of the core metabolic reconstruction initially had no gene attributed with producing the enzyme responsible for its catalysis. Similarly, many genes with no clear biological function were found to be essential in the Tn-seq screen. By attempting to fill the gaps in the *in silico* model with the uncharacterized essential genes, we were able to assign putative functions to eight previously uncharacterized genes (Table S4). Two of these genes were chosen for further characterization, *smc01361* and *smc04042.* The *smc01361* gene was annotated as encoding a dihydroorotase, and mutation of *smc01361* resulted in pyrimidine auxotrophy (Figure S14). Given its location next to *pyrB,* and the presence of an essential PyrC dihydroorotoase encoded elsewhere in the genome (Data Set S1), we propose that *smc01361* encodes an inactive dihydroorotase (PyrX) required for PyrB activity as has been observed in some other species including *Pseudomonas putida* [35,36]. The essential *smc04042* gene was annotated as an inositol-1-monphosphatase family protein. It was previously observed that rhizobia lack a gene encoding a classical L-histidinol-phosphate phosphohydrolase, and it was suggested an inositol monophosphatase family protein may fulfill this function instead [37]. Mutation of *smc04042* resulted in histidine auxotrophy (Figure S14), consistent with this enzyme fulfilling the role of a L-histidinol-phosphate phosphohydrolase. It is likely that this is true for most rhizobia, as putative orthologs of this gene were identified in all 10 of the examined Rhizobiales genomes (Data Set S4). These examples illustrate the power of the combined Tn-seq and metabolic reconstruction process in the functional annotation of bacterial genomes.

## DISCUSSION

In this study, we developed a new variant of the Tn5 transposon for construction of non-polar insertion mutations that should be readily adaptable for use with other α-proteobacteria. The Tn5 transposon was chosen as it was expected to have low insertion site specificity in *S. meliloti* [38]. However, we observed a moderate sequence insertion bias for GC rich regions consistent with previous studies of the Tn5 transposon [39–41]. The consensus sequence of ∼ 190,000 unique insertion locations was largely consistent, but not identical, with that previously reported [41]; however, the specificity appeared to extend past the 9 base pair region that is duplicated during Tn5 insertion (Figure S3). While this bias is unlikely to have a significant influence on the results in species with high GC content genomes, such as *S. meliloti,* accounting for this bias may be important when applying Tn5 mutagenesis to species with low GC content genomes.

Greater than 10% of species with a sequenced genome contain a genomic architecture similar to *S. meliloti,* that is, with at least two large DNA replicons [42,43]. Several studies have revealed that, in many ways, each replicon acts as a functionally and evolutionarily distinct entity (for a review, refer to [43]); yet, there can also be regulatory cross-talk [44], as well as the exchange of genetic material between the replicons [45]. The Tn-seq analyses reported here provide new insights into the functional integration of secondary replicons into the host organism. The pSymB replicon of *S. meliloti* is known to have two essential genes (which were transferred to the chromosome for this study) [45], while pSymA has no essential genes [46]. However, the large number of chromosomal genes—across many functional groups (Figure 2)— that became conditionally essential following the removal of pSymA and pSymB indicate the presence of many genes whose products can perform essential metabolic capabilities but that remain cryptic due to inter-replicon epistatic interactions. It was also interesting to note that the strength of the correlation between duplicates (Figure S5), as determined by the size of the absolute residuals, was higher for the ΔpSymAB strain than for the wild type strain in both media (p-value < 2.2 × 10^−16^ for both media, as determined with Welch two-sample t-tests). This may be reflective of the genetic robustness encoded by the secondary replicons and the stochastic activation of these processes in the mutant population. Potentially, the high level of inter-replicon redundancy may reduce the level of purifying selection on the chromosomal copies of the genes, facilitating more rapid diversification of gene functionality and increased rates of chromosomal gene evolution. Overall, the results of these analyses suggest that secondary replicons may influence the evolution of the chromosome and play a vital role in the biology of the organism, even if these activities remain cryptic due to inter-replicon epistatic interactions.

More generally, the Tn-seq data reported here provide a unique perspective of how a gene’s genomic environment influences its genotype-phenotype relationship. Previous studies have illustrated that the fitness phenotypes of orthologous genes of both distant and closely related species may differ [21,23–28,47], and even how intercellular effects within microbial communities can modify the essential genome of a species [48]. The data reported here more directly addressed the influence of the genomic environment by comparing the fitness phenotypes of mutating the exact same set of ∼ 3,500 genes in two very different genomic environments. It was found that the non-essential genome had a remarkable influence on what was classified as a growth-promoting gene, with 10% of *S. meliloti* chromosomal genes exhibiting fitness-based genetic interactions with the non-essential component of the genome (Figure 2). This observation was not growth medium-dependent, was not unique to a specific gene functional class, and was not simply due to an overall reduced fitness of the ΔpSymAB strain as the findings could be largely replicated *in silico* (Figure 4).

The majority of the genes whose fitness phenotype was dependent on the genomic environment became more important for fitness following the genome reduction. In many cases, this is expected to reflect a loss of functional redundancy; the increased importance of the chromosomal cytochrome genes likely reflects a compensation for the loss of the pSymA/pSymB encoded cytochrome complexes (Figure 6). In other cases, it may reflect newly activated pathways that must compensate for the loss of a normal housekeeping pathway. The specific essentiality of proline biosynthesis, and the second half of histidine biosynthesis, in the ΔpSymAB strain during growth in rich medium presumably reflects the inability of these strains to transport these compounds and must therefore synthesize them *de novo* (Figure 6). Indeed, previous metabolomics work is consistent with the ΔpSymAB strain being unable to transport many amino acids, including proline and histidine [49]. Similarly, glycolysis appeared specifically essential in the ΔpSymAB strain in rich medium (Figure 6), likely as the reduced metabolic capacity of this strain [29] led to a greater reliance on catabolism of the abundant sucrose for energy and biosynthetic precursors. Specific gene essentiality in the ΔpSymAB background may also occur as a result of synthetic negative interactions that are not associated with metabolic redundancy, for example, synthetic effects of disrupting two independent aspects of the cell envelope. This may be reflected in the specific essentiality of the *feuNPQ* and *ndvAB* genes involved in production of periplasmic cyclic β-glucans (Figure 6) [50–53]. The cell envelope of the ΔpSymAB strain is altered compared to the wild-type, due to the loss of succinoglycan production [54] and the *bacA* gene [55], and the membrane lipid composition contains signs of increased stress [49]. The fitness of disrupting periplasmic cyclic β-glucans biosynthesis in this background, further altering the cell envelope, may therefore represent a synthetic negative interaction.

**Figure 6.**
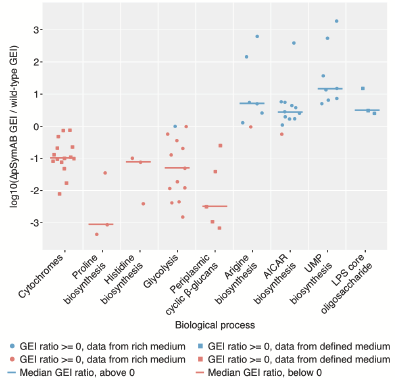
Gene essentiality index (GEI) changes for genes of selected biological pathways. Each circle or square represents an individual gene, and shows the logio of the ratio of the GEI for that gene in the ΔpSymAB background compared to the wild-type background. Lines indicate the median value of all genes included from the biological process. The underlying data is given in Table S9. Genes included in each process are as follows: Cytochrome C oxidase related genes -*ctaB, ctaC, ctaD, ctaE, ctaG, ccsA, cycH, cycJ, cycK, cycL, ccmA, ccmB, ccmC, ccmD, ccmG;* Proline biosynthesis -*proA, proBl, proC;* Histidine biosynthesis -*hisB, hisD, smc04042*; Glycolysis and related genes -*glk, frk, pgi, zwf, pgl, edd, eda2, gap, pgk, gpmA, eno, pykA, pyc;* Periplamic cyclic P-glucan biosynthesis -*feuN, feuP, feuQ, ndvA, ndvB;* Arginine biosynthesis -*argB, argC, argD, argF1, argG, argH1, argJ;* AICAR biosynthesis -*purB, purC, purD, purE, purF, purH, purK, purL, purM, purN, purQ, smc00494;* UMP biosynthesis -*carA, carB, pyrB, pyrC, pyrD, pyrE, pyrF, smc01361;* LPS core oligosaccharide biosynthesis -*lpsC, lpsD, lpsE.*

Somewhat surprisingly, approximately a quarter of the genes with a genomic environment effect had a greater fitness defect in the wild type strain. In some cases this may have been due to the reduced nutrient demand of the ΔpSymAB strain as a result of the smaller genome content. For example, mutations of genes for arginine biosynthesis and the biosynthesis of AICAR and UMP, common precursors in the synthesis of purines and pyrimidines, respectively, had fitness defects in rich medium specifically in the wild-type (Figure 6). This may reflect that in this environment, the uptake of these nutrients is growth limiting to the wild-type in the absence of their *de novo* synthesis, whereas this is not the case in the ΔpSymAB strain due to the reduced genome size, and thus lower nutrient requirement, and the already reduced growth rate (Figure S8). Another possibility is that removal of pSymAB evokes phenotypes that are epistatic to many of those brought about by chromosomal mutations. For example, the removal of pSymB is expected to have resulted in alterations of the cell membrane [49,54,55]; our observation that many mutations causing greater relative fitness defects in wild-type cells are associated with lipid metabolism, such as biosynthesis of the lipopolysaccharide core oligosaccharide (Figure 6) may be a result of those mutations being phenotypically masked in the absence of pSymB.

Our work in integrating the Tn-seq data with *in silico* metabolic modelling made it evident that Tn-seq alone is insufficient to identify the entire core metabolism of an organism; almost a third of the reactions present in the core metabolic reconstruction were not supported by Tn-seq data (Figure 5 and Table 3). Similarly, the large number of changes made in the gene-reaction relationships when producing the core model illustrated the limitations in the quality of metabolic reconstructions when high-throughput mutagenesis data are lacking. In some cases, the gaps in the Tn-seq data were due to genomic environment effects, such as genetic redundancy, in other cases it was due to the inclusion of reactions that are non-essential but that are nonetheless required for production of ‘wild type’ cells, and sometimes the gene associated with a reaction is simply unknown. A fourth possibility is phenotypic complementation through cross-feeding. Given that Tn-seq involves growth of a population of mutants, a mutant unable to produce an essential metabolite may still grow if the metabolite is excreted and transferred to the mutant from the rest of the population.

Regardless of the reasons why Tn-seq may have missed so many central metabolic reactions, this limitation can have a significant practical impact in the modern era of synthetic biology. The results of Tn-seq studies may be used to guide engineering of designer microbial factories with specific properties [56], or for the identification of putative new therapeutic targets [25,57]. While Tn-seq studies undoubtedly give invaluable information to be used towards these goals, basing engineered cells solely on Tn-seq studies is insufficient, as evidenced in the recent monumental efforts to design and synthesize a minimal bacterial genome [22]. Importantly, this limitation can be overcome by combining Tn-seq with metabolic modelling. We are aware of only a few other studies making use of both Tn-seq data and metabolic reconstruction [58–62]; however, these studies almost always focus on using the Tn-seq data to refine the metabolic reconstruction. As illustrated here, combining an experimental Tn-seq approach with a ground-up *in silico* metabolic reconstruction strategy can improve not only the reconstruction but also overcome the limitations of the Tn-seq approach. A Tn-seq-guided reconstruction process forces the identification of missing essential reactions, while ensuring correct gene-reaction associations, and the integrated approach can facilitate functional annotation of genes without clear biological roles. This process allows one to obtain a very high-quality representation of the metabolism, and the underlying genetics, of the organism in the given environment. The resulting model can serve as a blueprint to simply understand the workings of the cell, or as a basis for developing new cell factories.

## MATERIALS AND METHODS

### Bacterial strains, media, and growth conditions

The wild type and ΔpSymAB strains used throughout this work are the RmP3499 and RmP3496 strains, respectively, whose construction was described previously [30]. All *E. coli* or *S. meliloti* strains used in this study are described in Table S5 and were grown at 37°C or 30°C, respectively. BRM medium was used as the rich medium for growth of the *S. meliloti* strains, and it consisted of 5 g/L Bacto Tryptone, 5 g/L Bacto Yeast Extract, 50 mM NaCl, 2 mM MgSO_4_, 2 μM CoCl_2_, 0.5% (w/v) sucrose, and supplemented with the following antibiotics, as appropriate: streptomycin (Sm, 200 μg/ml), neomycin (Nm, 100 μg/ml), gentamycin (Gm, 15 μg/ml). The defined medium for growth of *S. meliloti* contained 50 mM NaCl, 10 mM KH_2_PO_4_, 10 mM NH_4_Cl, 2 mM MgSO_4_, 0.2 mM CaCl_2_, 0.5% (w/v) sucrose, 2.5 μM thiamine, 2 μM biotin, 10 μM EDTA, 10 μM FeSO4, 3 μM MnSO4, 2 μM ZnSO4, 2 μM H3BO3, 1 μM CoCl_2_, 0.2 μM Na_2_MoO_4_, 0.3 μM CuSO_4_, 50 μg/ml streptomycin, and 30 μg/ml neomycin. *E. coli* strains were grown on Luria-Bertani (LB) supplemented with the following antibiotics as appropriate: chloramphenicol (30 mg/ml), kanamycin (Km, 30 μg/ml), gentamycin (Gm, 3 μg/ml).

### Growth curves

Overnight cultures grown in rich media with the appropriate antibiotics were pelleted, washed with a phosphate buffer (20 mM KH_2_PO_4_ and 100 mM NaCl), and resuspended to an OD600 of 0.25. Twelve μl of each cell suspension was mixed with 288 μl of growth medium, without antibiotics, in wells of a 100-well Honeycomb microplate. Plates were incubated in a Bioscreen C analyzer at 30°C with shaking, and OD600 recorded every hour for at least 48 hours.

### *S. meliloti* mutant construction for Tn-seq validation

Single gene knockout mutants were generated through single cross-over plasmid integration of the suicide plasmid pJG194 [63]. Approximately 400-bp fragments homologous to the central portion of the target genes were PCR amplified using the primers listed in Table S6. PCR products as well as the pJG194 and pJG796 vectors were digested with the restriction enzymes *Eco*RI/*Hind*III, *Bam*HI/*Xba*I, or *Sal*I/*Xho*I, and each PCR fragment was ligated into the appropriately digested pJG194 or pJG796 vector using standard molecular biology techniques [64], and all recombinant plasmids verified. Recombinant plasmids were mobilized from *E. coli* to *S. meliloti* via tri-parental matings as described before [52], and transconjugants isolated on BRM Sm Nm agar plates. All *S. meliloti* mutants were verified by PCR.

Transduction of the integrated plasmids into the *S. meliloti* wild type and ΔpSymAB strains was performed using phage N3 as described elsewhere [65], with transductants recovered on BRM medium containing the appropriate antibiotics.

### Construction of the transposon delivery vector pJG714

The plasmid pJG714 is a variant of the previously reported mini-Tn5 delivery plasmid, pJG110 [63], with the primary modifications being removal of the *bla* gene and pUC origin of replication, and introduction of the *pir*-dependent R6K replication origin. A map of pJG714 is given in Figure S1A, and the complete sequence of the transposable region is provided in Figure S1B. This delivery plasmid is maintained in *E. coli* strain MFD*pir* [66], which possesses chromosomal copies of R6K *pir* and RK2 transfer functions. MFDpir is unable to synthesize diaminopimelic acid (DAP), thus disabling growth on rich or defined medium lacking supplemental DAP. The MFD*pir*/pJG714 strain is cultured on rich medium containing kanamycin and 12.5 μg/ml DAP.

### Tn-seq experimental setup

Transposon mutagenesis was accomplished in the wild-type and ΔpSymAB strains in parallel. Flask cultures of MFDpir/pJG714 and the two *S. meliloti* strains were grown overnight to saturation, and pellets were washed and suspended in BRM to a final OD600 value of approximately 40. Equal volumes of each suspension were mixed as bi-parental matings, to accomplish mobilization of the transposon delivery vector into the *S. meliloti* recipient strains. These cell mixtures were plated on BRM supplemented with 50 μg/ml DAP and incubated at 30°C for 6 h. Mating mixtures were collected in BRM with 10% glycerol, and cell clumps were broken up by shaking the suspended material for 30 min at 225 rpm. Aliquots were stored at −80°C. For selection of transposants, mating mixes were thawed and plated at a density of 15,000 cfu/plate (150-mm plates) on BRM supplemented with Sm and Nm. To accomplish equivalent coverage of each genome with transposon insertions, 675,000 and 360,000 colonies were selected for the wild-type and ΔpSymAB strains, respectively. For each recipient, transposon mutant colonies were collected and cell clumps were broken up as described above. The selected clone libraries were aliquoted and stored at −80°C.

For whole-population selection and massively parallel sequencing of transposon ends, 1×10^9^ cells from each of the two clone libraries were transferred into 500 ml of either BRM or defined medium, allowing approximately 8-10 generations of growth at 30°C before reaching saturation. At this stage, cells were pelleted, DNA was extracted using the MoBio microbial DNA isolation kit (*#*12255-50), and the resulting DNA was fragmented with NEB fragmentase (#M0348S) to an average molecular weight of 1000 bp. After clean-up (Qiagen #27106), the resulting DNA fragments were appended with short 3’ homopolymer (oligo-dCTP) tails using terminal deoxynucleotidyl transferase (NEB #M0315S), and this sample was used as the template for a two-round PCR that gave rise to the final Illumina-ready libraries. In the first round, a transposon end-specific primer (1TN) and oligo-G primer (1GG) were used (all primer sequences can be found in Table S6). After clean-up, a portion of the first-round product was used as the template for the second-round reaction employing a nested transposon-specific primer (2TNA-C) and a reverse index-incorporating primer (2BAR01-08). The series of three 2TN primers (A–C) were designed to incorporate base diversity in the opening cycles of Illumina sequencing, and the series of eight 2BAR primers were designed to uniquely identify each experimental condition in a single multiplexed sequencing sample. After PCR amplification of transposon-flanking sequences with concomitant incorporation of Illumina adapters and barcodes, the samples were size-selected for 200-600-bp fragments, and sequenced on an Illumina Hi-Seq instrument as 50-bp single-end reads. Raw reads were used as input into a custom-built Tn-seq analytical pipeline, which was recently described [57].

### Calculation of gene and synthetic indexes

For calculation of Gene Essentiality Index (GEI) scores, a pseudo count of one was first added to all gene read counts for each replicate. GEI were then calculated by summing the number of reads that mapped to the gene in both replicates, and dividing this number by the nucleotide length of the gene. GEI scores were calculated for each gene separately in each medium and in each strain. All GEI values are available in Data Set S1.

Synthetic Media Index (SMI) scores were calculated to represent the difference in GEI scores between the two media for the same strain. Raw SMI scores were determined by dividing the GEI of the gene in defined medium by the GEI of the gene in rich medium. Processed SMI scores, those shown throughout the manuscript, were determined as follows. If the raw value was above one, the processed SMI and the raw SMI are the same. Raw SMI scores that were below one were converted to processed SMI scores through the transformation, “1 / raw SMI score”, and presenting the value as a negative number.

Raw and processed Synthetic Rich Index (SRI) and Synthetic Defined Index (SDI) scores were calculated to represent the difference in the GEI scores of a gene between the wild-type and ΔpSymAB strains when grown in rich or defined medium, respectively. SRI and SDI indexes were calculated using the same procedure as described for the SMI scores above. All synthetic index scores are provided in Data Set S1.

### Statistical analysis of the Tn-seq output

The output of the Tn-seq analysis pipeline was used in the fitness classification of genes as follows. First, all genes with no observed insertions were classified as essential. Next, GEI scores were imported into R version 3.2.3 and log transformed. Initial clustering of the log transformed GEI scores into fitness categories was performed using the *Mclust* function of the *Mclust* package in R [67]. In short, this function attempts to explain the distribution of GEI values by fitting a series of overlapping Guassian distributions, with the number and shape of the distributions determined by *Mclust.* The data are then assigned to different categories based on the probability of the data point arising from each of the distributions. As high uncertainty in the classification of genes at the borders of groupings exists, the clusters were refined through the use of affinity propagation implemented by the *apcluster* function of the *apcluster* package of R [68]. All genes belonging to an *apcluster* grouping that contained an essential gene, as determined in any of the previous steps, were re-annotated as essential. Additionally, all genes belonging to an *apcluster* grouping that spanned the border of two *Mclust* goups were transferred to the same classification, based on which cluster the genes had a higher median probability of being derived from in the *Mclust* analysis. Finally, genes that were classified as ‘essential’ in one medium and ‘large growth impairment’ in the second medium, but that were identified as having no medium specificity based on their SMI scores, were considered as essential in both media.

Genes with GEI scores significantly different between conditions were determined as follows. The synthetic indexes (SMI, SDI, SRI) scores were imported into R and log transformed, and the following clustering performed independently for each index. The log transformed synthetic scores were clustered using *Mclust* and *apcluster* in R as described above for the GEI scores. In the case of the SMI scores, three clusters were produced ‘Little to no difference’, ‘Moderate difference’, and ‘Large difference’; only genes with a SMI scores classified as ‘Large difference’ were considered to display a medium specificity. In the case of SDI and SRI scores, only two clusters were produced: ‘Little to no difference’ and ‘Difference between strains’.

### Gene functional enrichments

Assignment of chromosomal genes into specific functional categories was performed largely based on the annotations provided on the *S. meliloti* Rm1021 online genome database (https://iant.toulouse.inra.fr/bacteria/annotation/cgi/rhime.cgi). This website pulls annotations from several databases including PubMed, Swissprot, trEMBL, and Interpro. Additionally, it provides enzyme codes, PubMed IDs, functional classifications, and suggested Gene Ontology (GO) terms for most genes. The numerous classifications were simplified to 18 functional categories, designed to adequately cover all core cellular processes. Occasionally, ambiguous or conflicting annotations were observed. In these cases, protein BLASTp searches through the NCBI server were performed against the non-redundant protein database. If putative domains were detected within the amino acid sequence, a combination of the best hit (lowest E-value) and consensus among domain annotations were used to categorize the gene in question. If no putative domains were detected, the functional annotation was based on the best scoring protein hits in the database. The functional annotations of all chromosomal genes are provided in Data Set S5.

### Data visualization

Tn-seq results were visualized using the Integrative Genomics Viewer v2.3.97 [69]. Scatter plots, functional enrichment plots, box plots, and line plots were generated in R using the *ggplot2* package [70]. Venn diagrams were produced in R using the *VennDiagram* package [71]. The genome map was prepared using the circos v0.67-7 software [72]; the sliding window insertion density was calculated with the *geom_histogram* function of *ggplot2,* and the GC skew was calculated using the analysis of sequence heterogeneity sliding window plots online webserver [73]. The metabolic model was visualized using the iPath v2.0 webserver [74]. The logo of the transposon insertion site specificity was generated by first extracting the nucleotides surrounding all unique insertion sites in one replicate of the wild-type grown in rich medium using Perl v5.18.2, followed by generation of a hidden Markov model with the *hmmbuild* function of HMMER v3.1b2 [75] and visualization with the Skylign webserver [76].

### Blast Bidirectional Best Hit (Blast-BBH) strategy

Putative orthologous proteins between species were identified with a Blast-BBH approach, implemented using a modified version of our in-house Shell and Perl pipeline [77]. This pipeline involved GNU bash v4.3.48(1), Perl v5.22.1, Python v2.7.12 and the Blast v2.6.0+ software [78]. Proteomes were downloaded from the National Center for Biotechnology Information repository, and the Genbank annotations were used. As a threshold to limit false positives, Blast-BBH pairs were only maintained if they displayed a minimum of 30% amino acid identify over at least 60% of the protein. To identify putative duplicate proteins in *S. meliloti,* the same Blast-BBH approach was employed to compare the *S. meliloti* chromosomal proteome with the proteins encoded by pSymA and pSymB.

### *In silico* metabolic modeling procedures

All simulations were performed in Matlab 2017a (Mathworks) with scripts from the Cobra Toolbox (downloaded May 12, 2017 from the openCOBRA repository) [79], and using the Gurobi 7.0.2 solver (www.gurobi.com), the SBMLToolbox 4.1.0 [80], and libSBML 5.15.0 [81]. Boundary conditions for simulation of the defined medium are given in Table S7. *In silico* analysis of redundancy in the *S. meliloti* genome was performed using the iGD1575b metabolic reconstruction, whose development is described in the following section. Single and double gene deletion analyses were performed using the *singleGeneDeletion* and *doubleGeneDeletion* functions, respectively, using the Minimization of Metabolic Adjustment (MOMA) method. All Matlab scripts used in this work are provided as File S3.

For all deletion mutants, the growth rate ratio (grRatio) was calculated (growth rate of mutant / growth rate of wild type). Single gene deletion mutants were considered to have a growth defect if the grRatio was < 0.9. For the double gene deletion analysis, if the grRatio of the double mutant was less than 0.9 the expected grRatio (based on multiplying the grRatio of the two corresponding single mutants), the double deletion was said to have a synthetic negative phenotype.

### Development of iGD1575b

For *in silico* analysis of redundancy in the *S. meliloti* genome, the previously published *S. meliloti* genome-scale metabolic model iGD1575 [34] was modified slightly. As indicated in Table S8, the biomass composition was updated to include 31 additional compounds at trace concentrations, including vitamins, coenzymes, and ions, in order to ensure the corresponding transport or biosynthetic pathways were essential. However, the original model iGD1575 was unable to produce vitamin B12 and holo-carboxylate. To rectify this, the reversibility of rxn00792_c0 was changed from ‘false’ to ‘true’, and the reactions rxn01609, rxn06864, and rxnBluB were added to the model. However, no new genes were included in the model. This updated model was termed iGD1575b and is available in SBML and Matlab format in File S2.

### Simulating the removal of pSymA and pSymB *in silico.*

Several modifications to iGD1575b were required in order to produce a viable model following the deletion of all pSymA and pSymB genes. As described previously [34], succinoglycan was removed from the biomass composition, ‘gapfill’ GPRs (gene-protein-reaction relationships) were added to the reactions ‘rxn01675_c0’, ‘rxn01997_c0’, ‘rxn02000_c0’, and ‘rxn02003_c0’ in order to allow the continued production of the full LPS molecule, as well as to ‘rxn00416_c0’ to allow asparagine biosynthesis. Additionally, ‘gapfill’ GPRs were added to the reactions ‘rxn03975_c0’ and ‘rxn03393_c0’ so that removal of pSymA and pSymB did not prevent biosynthesis of vitamin B12 and ubiquinone-8, respectively. Finally, a glycerol export reaction via diffusion (rxnBLTPcpd00100b) was added to remove the glycerol build-up resulting from cardiolipin biosynthesis. The modified version of the model was termed iGD1575c, and is available in in SBML and Matlab format in File S2. For simulating the removal of pSymA and pSymB in Matlab, all pSymA and pSymB genes were deleted from the iGD1575b model using the *deleteModelGenes* function, followed by the removal of all constrained reactions using the *removeRxns* function.

### Building the draft *R. leguminosarum* metabolic model

A draft, fully automated model containing no manual curation for *R. leguminosarum* bv. *viciae* 3841 was built using the KBase webserver (www.kbase.us). The Genbank file (GCA_000009265.1_ASM926v1_genomic.gbff) of the *R. leguminosarum* genome [82] was uploaded to KBase and re-annotated using the ‘annotate microbial genome’ function, maintaining the original locus tags. An automated metabolic model was then built using the ‘build metabolic model’ function, with gap-filling. This model included 1537 genes, 1647 reaction, and 1731 metabolites, and is available in in SBML and Matlab format in File S2. The biomass composition was not modified from the default Gram negative biomass of Kbase. All essential model genes were determined using the Cobra Toolbox in Matlab with the *singleGeneDeletion* function and the MOMA protocol, with exchange reaction bounds set as provided in Table S7.

### Building the *S. meliloti* core metabolic reconstruction, iGD726

The iGD726 model was built from the ground-up using the existing iGD1575 model as a reaction and GPR database, and with the Tn-seq data as a guide. Each metabolic pathway included in iGD726 was rebuilt in a new file by adding individual reactions to the file. These reactions were taken from iGD1575, or were taken from other sources, primarily the Kyoto Encyclopedia of Genes and Genomes (KEGG) database [83], if an appropriate reaction was missing in iGD1575. Following the transfer of each reaction, the genes associated with the reaction were checked against the Tn-seq data, and a literature search for each associated gene was performed. The gene associations were then modified as necessary to ensure the model accurately captured the experimental data. For example, if gene was experimentally determined to be essential, but the corresponding reaction for the gene was associated with multiple alternative genes, all but the essential gene were removed from the reaction. Similarly, if a non-essential gene was associated with an essential reaction, a second gene or an Unknown was added to reflect the apparent redundancy in the genome. Where possible, unknowns in the gene associations were replaced with genes whose gene product may catalyze the reaction.

During the construction of the core model, the biomass composition was updated. This included modifying the membrane lipid composition to include lipids with different sized fatty acids based on the ratio experimentally determined [84]; the original iGD1575 model contained only one representative per each membrane lipid class. Additionally, essential vitamins, cofactors, and ions were added to the biomass composition at trace concentrations to ensure that their biosynthesis or transport was essential. The complete biomass composition is provided in Table S3.

The necessary metabolic and transport reactions to allow the model to growth with sucrose, glucose, or succinate were included in the reconstruction. Once the model was capable of producing all biomass components using any of the three carbon sources, the list of model genes was compared with the list of 489 core growth promoting genes to identify genes not included in the model but experimentally determined to contribute to growth. When possible, missing genes and their corresponding reactions were added to the core model. The final model contained 726 genes, 681 reactions, and 703 metabolites, and is provided in SBML and Matlab format in File S2, and as an Excel file in Data Set S4. The Excel file contains all necessary information for use as a *S. meliloti* metabolic resource, including the reaction name, the reaction equation using the real metabolite names, the associated genes/proteins, and references. Additionally, for each reaction, the putative orthologs of the associated genes in 10 related Rhizobiales species are included, allowing the model to provide useful information for each of these organisms.

## FUNDING INFORMATION

This work was funded by National Science Foundation grant IOS-1054980 to Joel S Griffitts. George C diCenzo was supported by the Natural Sciences and Engineering Research Council of Canada (NSERC) through a PDF fellowship. Work in the laboratory of Turlough M Finan was funded by NSERC. The funders had no role in the study design, data collection and interpretation, or the decision to submit the work for publication.

